# A Mechanism Driving Echinocandin Heteroresistance in *Candida glabrata*

**DOI:** 10.64898/2026.04.16.718912

**Authors:** Abigail A. Harrington, Kyle W. Cunningham

## Abstract

While pathogenic fungi can acquire resistance to the current arsenal of antifungals through genetic mutations, heteroresistance has emerged as an important new cause of therapeutic failures. Heteroresistance is generally thought to arise in small subpopulations that display phenotypic resistance to antifungals without genetic mutations. This study blurs that line by showing gain-of-resistance mutations in *FKS2*, which encodes a target of echinocandins and fungerps, cause large amounts of heteroresistance in *Candida glabrata* through heterogeneous expression of the gene even in clonal cell populations. Heteroresistance decreased when stress-responsive transcription factors (Crz1, Rlm1) were eliminated and was nearly abolished when the upstream regulators (calcineurin, Slt2) were mutated or inhibited. Identical gain-of-resistance mutations in *FKS1*, a paralog of *FKS2*, showed much less heteroresistance due to its constitutive expression coupled with variable levels of antagonism by wild-type *FKS2*. A genome-wide screen using Tn-seq revealed additional regulators of heteroresistance and resistance including *IRA1*, an inhibitor of the Ras1-PKA signaling pathway that senses glucose availability. *IRA1* increased expression of *FKS2* and decreased expression of *FKS1*, which increased heteroresistance and decreased resistance, respectively, when these genes carried resistance mutations. Similar principles may govern heteroresistance in other fungal pathogens such as *Candida parapsilosis*, which naturally carries resistance mutations in *FKS1* and frequently exhibits heteroresistance to echinocandins, and *Candidozyma auris*, which easily acquires such mutations.

## INTRODUCTION

*Candida glabrata* (a.k.a. *Nakaseomyces glabratus*) is an opportunistic fungal pathogen of increasing clinical concern. Status as the second most common causative agent of candidiasis, only behind *Candida albicans*, has led the World Health Organization to list *C. glabrata* as a high priority threat. Recent surveillance in the United States indicated that *C. glabrata* was the most common cause of candidemia-related hospitalizations in 2024 [1]. This is in part due to *C. glabrata*’s innate resistance to azoles and rapid acquisition of resistance to echinocandins [2-4] that result in high rates of therapeutic failure [5, 6]. Resistance refers to the ability of cells to grow in higher doses of drug and is typically the result of genetic gain-of-resistance mutations in the drug target, or mutations in other components that increase drug efflux, prevent drug influx, or otherwise sequester the drug such that the typical drug dose is no longer adequate for complete clearance. Echinocandins and fungerps target essential enzymes in the plasma membranes of fungal cells that are responsible for synthesis of beta-1,3-glucan. In *C. glabrata*, the catalytic subunits of the synthases are encoded by two functionally redundant genes (*FKS1* and *FKS2*) that originated from an ancient whole-genome duplication event [7] with a third homologous gene (*FKS3*) playing no detectable roll [8]. *FKS1* expression is thought to be constitutive and unresponsive to cell wall stresses, while *FKS2* expression is highly responsive to stresses, particularly through the calcineurin and cell wall integrity (CWI) signaling pathways [8-10]. Strong resistance to echinocandins usually arises through non-synonymous mutations in either *FKS1* or *FKS2* within two different “hotspot” regions [6, 11]. Many of these hotspot mutations confer cross-resistance to multiple echinocandins (caspofungin, micafungin, and anidulafungin) and fungerps (enfumafungin and ibrexafungerp) [11-13].

Recently, enfumafungin was found to bind directly in the hotspot-1 region of Fks1 and Fks2 enzymes from the model yeast, *Saccharomyces cerevisiae*, with critical interactions between the antifungal and the conserved amino acids F639 and F658, respectively [14]. Substitution or deletion of this critical amino acid conferred cross-resistance to all echinocandins in addition to the fungerps without strongly altering enzymatic activity *in vitro* [14]. Mutations at the same sites in *FKS1* and *FKS2* genes of *C. glabrata* (F625 and F659) confer similar cross-resistance and arise frequently in clinical isolates from treated patients [8, 15-17].

Heteroresistance refers to a phenomenon where only a small subpopulation of cells within a clonal population transiently exhibits phenotypic resistance to antifungals in the absence of stable *de novo* mutations in the genome. Though heteroresistance has been associated with therapeutic failures, the epigenetic mechanisms that govern the phenotype remain poorly defined in pathogenic yeasts. Heteroresistance to azole-class antifungals such as fluconazole has been observed frequently in clinical isolates of *Cryptococcus* and attributed to rare duplications of a whole chromosome expressing the drug target and a drug efflux gene [18-21]. Heteroresistance to fluconazole in *C. glabrata* was associated with increased drug efflux and difficult-to-treat infections *in vivo* [22]. However, heteroresistance to echinocandins has not been observed in several isolates of *C. glabrata* [22, 23] and not yet reported in *C. albicans* [24]. Importantly, several clinical isolates of *Candida parapsilosis* and *Candida auris* were recently shown to exhibit clear heteroresistance to echinocandins leading to breakthrough infections [25].

Resistance to echinocandins can be quantified accurately using growth assays, such as broth microdilution assays, E-tests, and halo assays, and the relevant parameter for comparison is the minimum inhibitory concentration (MIC) or the drug concentration causing 50% growth inhibition (IC50). Though growth assays can sometimes provide information on minor subpopulations, heteroresistance is quantified most accurately and directly by the population analysis profile (PAP) assay [26-28]. This method relies on serially diluting clonal populations of fungal cells, plating on agar media containing serial dilutions of antifungal drugs, counting visible colonies after an appropriate incubation period, and calculating the percentage of CFUs relative to the untreated population. Non-heteroresistant isolates exhibit nearly all-or-none behavior at transitional doses, with colony forming units (CFUs) near 100% or 0% relative to the untreated population. Heteroresistant isolates exhibit CFUs between 50% and 0.0015% (limit of detection) over a broad range of doses. In the case of *C. parapsilosis* exposed to echinocandins, some isolates exhibited gradual decline in CFUs within the transitional range of doses, while others exhibited step-like declines [25], suggesting complexity in the underlaying regulatory mechanisms.

This study shows for the first time that *C. glabrata* strains with gain-of-resistance (GOR) mutations in *FKS2*, but not *FKS1*, exhibit strong heteroresistance to echinocandins. The gradual decline of CFUs over the transitional doses suggested that the cause of progressive heteroresistance may be variable expression of the *FKS2-GOR* in the clonal populations, which we confirmed genetically by introducing mutations in the calcineurin-Crz1 and Slt2-Rlm1 signaling pathways. Chemical stresses that increase *FKS2-GOR* expression exacerbated heteroresistance, while chemically decreasing *FKS2* expression attenuated heteroresistance to the echinocandin micafungin. A genome-wide Tn-seq screen in the heteroresistant strain *FKS2-F659Δ* was performed to identify additional regulators of heteroresistance. *FKS1* and *FKS2* were found to be mutually antagonistic, while *IRA1*, predicted to encode an inhibitor of the small GTPase Ras1 [29], was found to promote heteroresistance in *FKS2-F659Δ*, but decrease resistance in *FKS1-F625Δ* through unknown mechanisms. These findings begin to unravel mechanisms of echinocandin heteroresistance in pathogenic yeasts and indicate the important role that calcineurin signaling may play in promoting a variety of survival mechanisms to antifungal assault.

## RESULTS

### FKS2 gain of resistance mutants exhibit progressive heteroresistance to echinocandins

Many clinical isolates of *C. glabrata* have been reported as heteroresistant to fluconazole [22], but heteroresistance to echinocandins has not been reported. Two commonly used clinical isolates were tested in population analysis profile (PAP) assays for heteroresistance: BG2, a vaginal isolate, and CBS138, a gastrointestinal isolate. As expected, both strains exhibited no heteroresistance to micafungin (Fig 1A) or other echinocandins. Loss-of-function mutations in *MRP20, ALG6*, and *PDR1* were previously shown to mildly alter resistance to echinocandins in the BG2 strain background [30]. Deletion mutants lacking these genes, as well as a *PDR1-GOF* (gain-of-function) mutant that increased echinocandin resistance [30], did not exhibit altered heteroresistance to micafungin (Fig. S1). These findings show that echinocandin resistance can be uncoupled from heteroresistance in *C. glabrata*.

**Figure 1.**
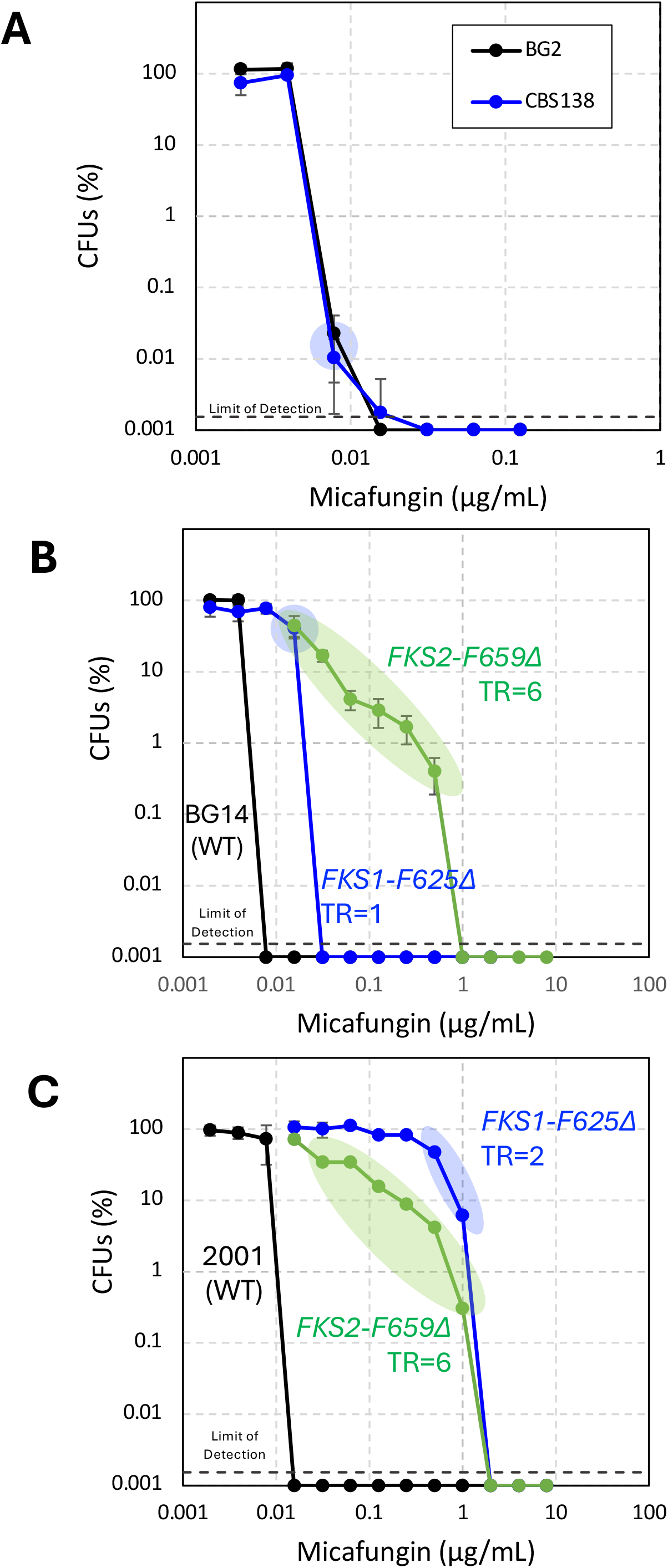
*C. glabrata* strains bearing *FKS2-F659Δ*, but not *FKS1-F625Δ*, exhibited strong progressive heteroresistance to micafungin. Wild-type clinical isolates (A) and derivatives carrying the same GOR mutation in *FKS1* and *FKS2* (B, C) were analyzed using the PAP assay. Replicate colonies were grown to saturation in SCD medium, serially diluted, and then plated on YPD agar media containing varying doses of micafungin. After 24 hours of incubation at 30°C, colonies were counted using a dissecting microscope. The percent CFUs for each strain at every dose were calculated with respect to the no drug control. The average CFUs of four biological replicates (±SD) is shown. Limit of detection = 0.0015%, or 1 colony detected out of 4 undiluted biological replicates.

Many isolates of *C. parapsilosis* exhibit heteroresistance to echinocandins [25]. As all isolates of this species bear a natural cross-resistance mutation in hotspot-1 of its *FKS1* gene [31], we tested whether hotspot-1 mutations in *FKS1* and *FKS2* genes of *C. glabrata* can confer heteroresistance in addition to resistance. Strikingly, the *FKS2-F659Δ* mutant derivative from strain BG2 exhibited broad heteroresistance to micafungin, whereas deletion of the same codon in *FKS1* (*FKS1-F625Δ*) exhibited only slight heteroresistance (Fig. 1B). The transitional range (TR) where CFUs varied from 50% maximum to the limit of detection was quantified by counting the doses as previously described [32]. The *FKS2-F659Δ* mutant exhibited TR=6 while the *FKS1-F625Δ* mutant exhibited TR=1, in comparison to the parent strain where TR=1. Similar results were found with the same mutants in 2001-HTU strain background (Fig. 1C), a well-studied derivative of CBS138 [33-35]. Similar results were also obtained using caspofungin in the agar medium instead of micafungin (Fig S2). Additionally, heteroresistance to micafungin was observed when several other gain-of-resistance substitution mutations were introduced into hotspot-1 of *FKS2*, but not when introduced into *FKS1*, albeit over a narrower range (Fig S3). Though these hotspot-1 mutations conferred elevated resistance to echinocandins in broth microdilution assays (Fig. S4) as described previously [11, 12, 36-38], only the *FKS2-GOR* mutants exhibited broad heteroresistance. The broadest heteroresistance was associated with *FKS2-F659Δ* mutants, in which a key amino acid residue required for binding of fungerps and echinocandins has been eliminated [8, 14-16]. The gradual decline of CFUs observed over six doses of micafungin is reminiscent of progressive heteroresistance in bacteria [39], suggesting that heterogeneous or variable expression of *FKS2-F659Δ* between cells may be responsible for heteroresistance in clonal populations.

### Calcineurin and CWI signaling pathways regulate heteroresistance

*FKS2* expression has been shown to be highly induced upon exposure to stresses, including echinocandin exposure [40, 41], *FKS1*-deficiency [42], and ER stresses [40, 43, 44]. *FKS2* expression in these conditions depends on several stress-activated signaling pathways such as the calcineurin signaling pathway [8] and the cell wall integrity (CWI) signaling pathway, at least in *S. cerevisiae* [42]. Previous work has shown that *FKS2* expression is variable cell-to-cell [10]. Differently, *FKS1* expression is unaffected by stress and is thought to be driven by a constitutive promoter [8]. To investigate the effects of calcineurin signaling on heteroresistance, the calcineurin dependent transcription factor, Crz1, was deleted in the *FKS2-F659Δ* strain background and tested. Interestingly, *FKS2-F659Δ crz1Δ* double mutant remained heteroresistant to micafungin (TR=7), but the number of CFUs were decreased an average of 80-fold (±SD) over the six transitional doses of micafungin (between 0.015625 and 0.25 µg/mL) (Fig 2A). Heteroresistance declined profoundly in strains when the calcineurin inhibitor, FK506 (2 µg/mL), was added to the agar media. However, FK506 had little or no effect on the CFUs observed in the *FKS1-F625Δ* mutant (Fig 2B). These findings suggest the progressive heteroresistance observed in *FKS2*-*GOR* strains of *C. glabrata* may be driven by variable levels of stress and stress signaling through calcineurin and Crz1 which influence *FKS2* expression. It is also possible that additional targets of calcineurin and Crz1 regulate Fks2 post-transcriptionally to promote heteroresistance.

**Figure 2.**
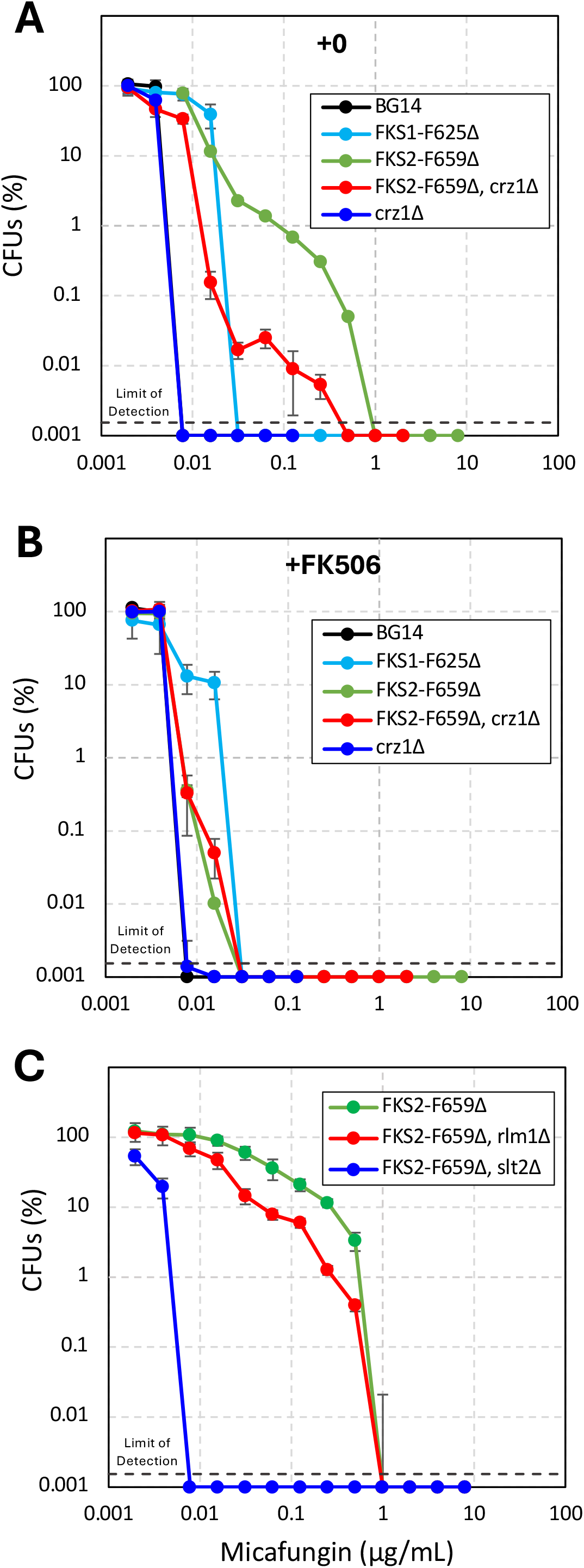
Calcineurin-Crz1 and Slt2-Rlm1 pathways promote heteroresistance in *FKS2-F659Δ* strains. PAP assays on the indicated were performed on the indicated derivatives of strain BG2 as described in Fig. 1 (A-C) or using agar medium supplemented with 2 µg/mL FK506 (B).

The effects of CWI signaling on heteroresistance were also investigated by deleting the genes encoding Slt2, a MAP-kinase, and a downstream transcription factor, Rlm1, in the *FKS2-F659Δ* strain background. The *FKS2-F659Δ rlm1Δ* double mutant maintained broad heteroresistance (TR=6) while exhibiting 5.3-fold decrease in CFUs compared to *FKS2-F659Δ* (TR=4) over the transitional range of micafungin (0.015625 and 0.5 µg/mL), indicating this transcription factor produces milder effects than Crz1 (Fig 2C). However, *FKS2-F659Δ slt2Δ* double mutants exhibited almost no heteroresistance (TR=1). The *rlm1Δ* and *slt2Δ* gene deletions were also generated in the wild-type parent strain BG14, a *ura3Δ* derivative of BG2, and the *FKS1-F625Δ* mutant backgrounds and tested for changes in heteroresistance and resistance. As expected, these CWI-deficient mutants were not detectably different from their respective parent strains (Fig S5). Together, these data suggest that major components of the CWI and calcineurin signaling pathways may be required for heteroresistance conferred by *FKS2-GOR* mutations. The milder roles of Rlm1 and Crz1 transcription factors suggest that other outputs of Slt2 and calcineurin may contribute to variable expression of *FKS2*-*GOR* in unstressed populations.

### Calcineurin regulates heteroresistance in subpopulations before exposure to echinocandins

The PAP assay counts colonies that arise from single cells plated on different doses of antifungals. To form a visible colony, the plated cell must survive the antifungal assault and proliferate along with most of its daughter cells. To distinguish whether calcineurin and stress alter heteroresistance of *FKS2*-*GOR* strains prior to plating, the *FKS2-F659Δ* mutant was exposed for just 2 hours to either 1 µg/mL FK506 or 0.6 µg/mL manogepix (MGX), an ER stressor that strongly activates of calcineurin and increases expression of *FKS2* [40] and then the cells were washed free of drugs and subjected to standard PAP assays. Predictably, the FK506 decreased CFUs 39-fold (averaged between 0.03125 and 0.25 µg/mL, TR=4), while MGX increased CFUs 5.1-fold (averaged between 0.0625 and 0.5 µg/mL, TR=4), both relative to the untreated *FKS2-F659Δ* strain (Fig 3). These results indicate that chemically decreasing or increasing calcineurin signaling in the population prior to plating directly influences heteroresistance of *FKS2-F659Δ* mutants through altered expression of *FKS2*. FK506 and MGX had no effect on heteroresistance of the wild-type parent strain (Fig 3).

**Figure 3.**
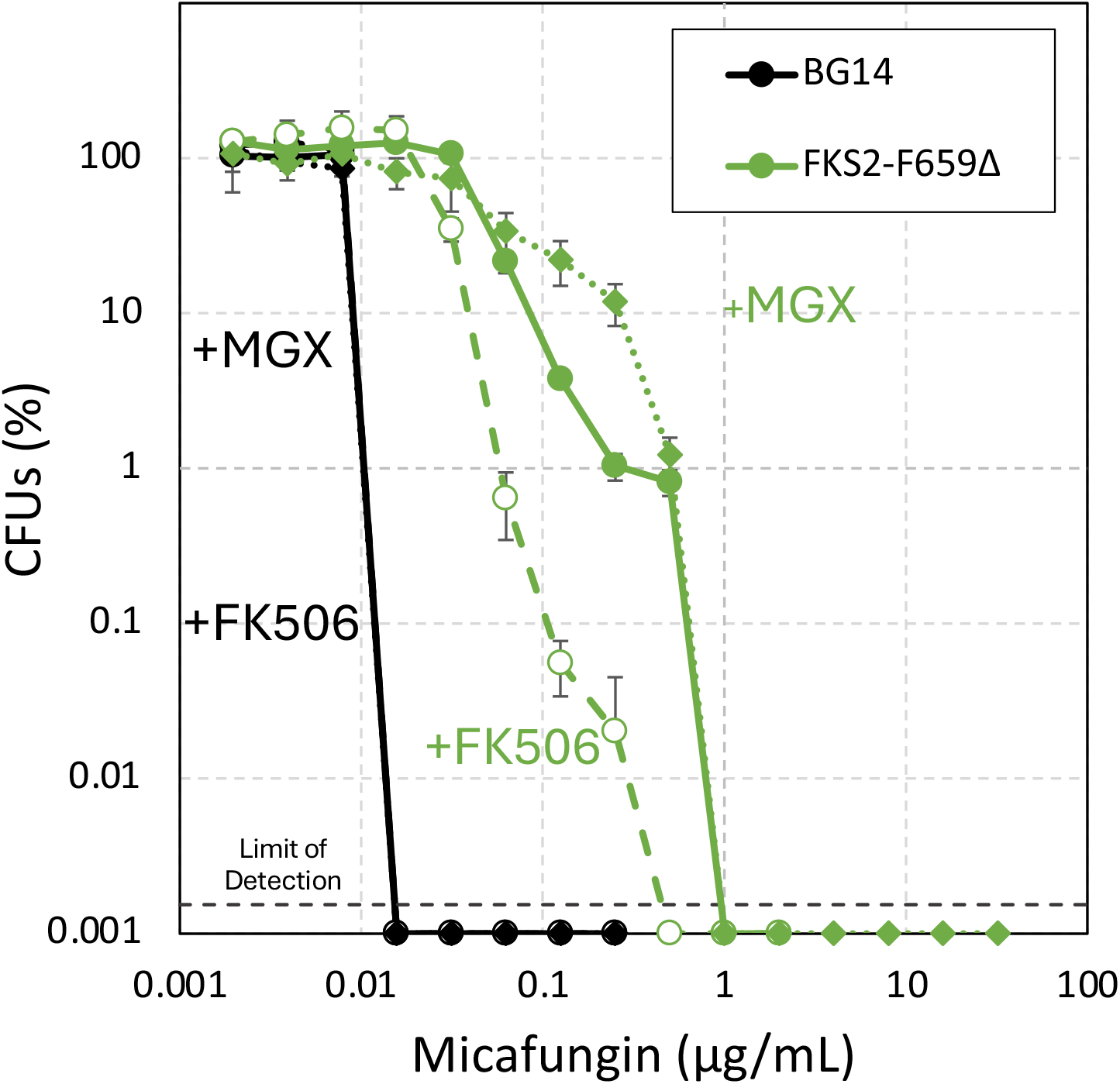
Increased and decreased calcineurin signaling caused increased and decreased heteroresistance, respectively. Derivatives of strain BG2, BG14 and *FKS2-F659Δ*, were grown to saturation, then diluted 1:50 into fresh SCD media containing either 1 µg/mL FK506, 0.6 µg/mL manogepix (MGX), or no drug and incubated at 30°C for 2 hours. The cells were collected, washed 1x in drug-free fresh medium, resuspended, and tested in PAP assays as described in Fig. 1.

### Tn-seq screens reveal additional regulators of echinocandin heteroresistance

Tn-seq quantifies the growth of many different transposon insertion mutants simultaneously during exposure to varying doses of antifungals and therefore may function well as a high-throughput PAP assay. To test this idea, a diverse pool of Hermes-NAT1 transposon insertions was generated in the *FKS2-F659Δ* mutant background, cultured in medium containing a broad range of caspofungin doses (0.25 to 32 µg/mL), and analyzed by Tn-seq [30]. Caspofungin was utilized in this experiment rather than micafungin to avoid the high number of reads in mitochondrial genes that confer micafungin resistance when disrupted [30]. The raw data were tabulated gene-wise and Z-scores for each gene were calculated as before [30] to identify specific gene disruptions that become significantly over- or under-represented relative to irrelevant genes. As expected, transposon insertions in *FKS2* produced strongly negative Z-scores (ranging from −14.7 to −6.5) at all eight doses of caspofungin due to the complete loss of resistance and heteroresistance conferred by the *FKS2-F659Δ* mutation (Fig. 4A). The *CRZ1* gene also exhibited mild under-representation at all doses of caspofungin (average Z = −2.0). *RLM1* was negative but not significant (average Z = −0.4), though *SMP1* (its paralog deriving from an ancient whole-genome duplication event) was significantly underrepresented (average Z = −3.0). *SMP1* has not been studied in *C. glabrata*, but functional redundance between Rlm1 and Smp1 might explain the mild loss of heteroresistance observed in *FKS2-F659Δ rlm1Δ* double mutant (Fig 2C).

**Figure 4.**
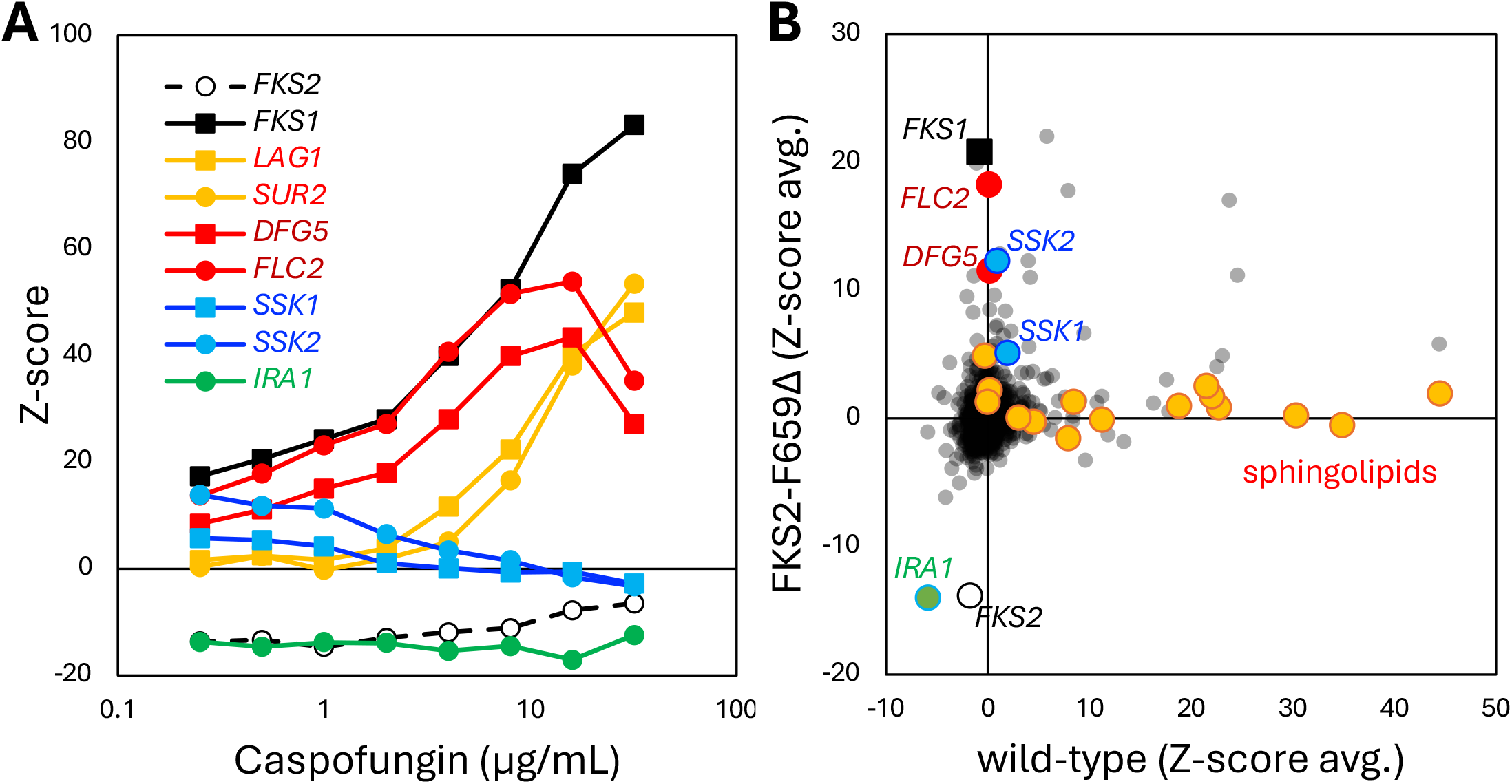
Tn-seq analysis of a *FKS2-F659Δ* mutant distinguishes regulators of heteroresistance and resistance. A diverse pool of *Hermes* transposon insertion mutants was generated in strain *FKS2-F659Δ*, divided, diluted into fresh SCD medium containing the indicated doses of caspofungin, incubated for 24 hr, and then analyzed by Tn-seq (see Methods) to identify genes with significant enrichment or depletion relative to the drug-free control cultures (positive or negative Z-scores). (A) Z-scores of selected genes are charted at each dose of caspofungin. (B) Z-scores of all genes in strain *FKS2-F659Δ* at the three lowest doses of caspofungin were averaged and charted against average Z-scores obtained previously from wild-type parent strain BG14 exposed to caspofungin [30]. Genes involved in sphingolipid biosynthesis (orange symbols) and selected other genes (colored symbols) are shown.

Earlier Tn-seq experiments using the isogenic wild-type parent strain identified twelve genes required for sphingolipid biosynthesis that could confer resistance to caspofungin when disrupted with transposons [30]. Seven of those genes, plus two additional ones (*AUR1, SCS7*), exhibited positive Z-scores in the *FKS2-F659Δ* mutant background, but the strongest effects were restricted to the four highest doses of caspofungin (44 to 32 µg/mL; Table S1). Two representative sphingolipid genes (*LAG1, SUR2*) were charted as a function of caspofungin dosage (Fig. 4A, orange symbols). Together, these findings show that *FKS2-F659Δ* was necessary for resistance at all doses of caspofungin and that sphingolipids were still required for inhibition of Fks2-F659Δ enzymes at the highest doses.

At the lowest doses of caspofungin, the Fks2-F659Δ enzyme was probably not directly inhibited due to its much weaker affinity for the drug [14]. Nevertheless, dozens of genes exhibited positive Z-scores at low doses indicating their disruption with transposons still improved fitness in these conditions, possibly due to heteroresistance rather than resistance. The Z-scores at three lowest doses of caspofungin were averaged for each gene in the *FKS2-F659Δ* dataset (heteroresistance) and charted against the average Z-scores from the wild-type parent strain obtained previously (resistance) (Fig. 4B). The correlation was poor (R^2^ = 0.051) and the sphingolipid genes clearly stood out as being more important in the wild-type parent strain than the *FKS2-F659Δ* mutant strain (orange symbols) at these doses of caspofungin. Conversely, several genes stood out with highly positive Z-scores in the *FKS2-F659Δ* mutant. Two genes (*DFG5, FLC2*) that may function together in attachment of GPI-anchored proteins to beta-1,3,-glucan in the cell wall [40] produced positive Z-scores in both strains (red symbols) and these Z-scores increased further at higher doses. Two genes that regulate the HOG signaling pathway (*SSK1, SSK2*; blue symbols) [45] exhibited positive Z-scores in *FKS2-F659Δ* at the lowest doses of caspofungin and then decreased at higher doses (Fig. 4A). The different patterns suggest complexity in the regulation of resistance and heteroresistance.

Strikingly, transposon insertions in *FKS1* produced strongly positive Z-scores at all caspofungin doses (Z = +17.4 to +83.2) in the *FKS2-F659Δ* mutant strain (Fig. 4A) while producing no such effect in the wild-type parent strain [30]. Thus, the drug-bound Fks1 protein somehow interfered with the performance of drug-resistant Fks2-F659Δ enzymes causing reduced fitness in caspofungin. Earlier studies have produced similar findings, where *FKS2-F659Δ fks1Δ* double mutants exhibited much higher resistance to echinocandins than *FKS2-F659Δ* single mutants [8, 17]. The converse relationship was also reported, where drug-bound Fks2-protein appeared to antagonize the activity of drug-free Fks1-F625Δ enzymes [8]. The molecular mechanisms responsible for this mutual antagonism between Fks and Fks-GOR proteins are not yet defined (see Discussion). The Tn-seq screen seemed to identify regulators of resistance, heteroresistance, and mutual antagonism that all impact the fitness conferred by Fks2-F659Δ.

### *IRA1* increases heteroresistance and decreases resistance

Genes required for heteroresistance of *FKS2-F659Δ* mutants should exhibit Z-scores similar to those of *FKS2* across all tested doses of caspofungin. A total of 30 genes exhibited average Z-scores less than −2.576, corresponding to p-value < 0.01, across all doses of caspofungin in the *FKS2-F659Δ* mutant (Table S1). GO term analysis (Table S2) [46] indicated this set was weakly enriched for components of the m-AAA complex (*YTA12, AFG3*) in the mitochondrial inner membrane that processes transit peptides. Further, two genes involved in chitin biosynthesis (*CHS3, CHS7*) were included in this set of 30 genes. By far, the strongest hit was *IRA1* (green symbols; Fig. 4A-B), which encodes an inhibitor of the small GTPase Ras1 that increases cAMP levels and signaling by protein kinase A in response to glucose availability in *S. cerevisiae* [29]. To test whether *IRA1* promotes heteroresistance, the gene was knocked out of *FKS2-F659Δ* mutants and wild-type strains and PAP assays were performed. Heteroresistance to micafungin was decreased 20-fold (averaged between 0.007813 and 0.5 µg/mL) in the *FKS2-F659Δ ira1Δ* double mutant (TR=8) relative to *FKS2-F659Δ* (Fig. 5A), consistent with a role for Ira1 in positively regulating expression or function of Fks2-F659Δ.

**Figure 5.**
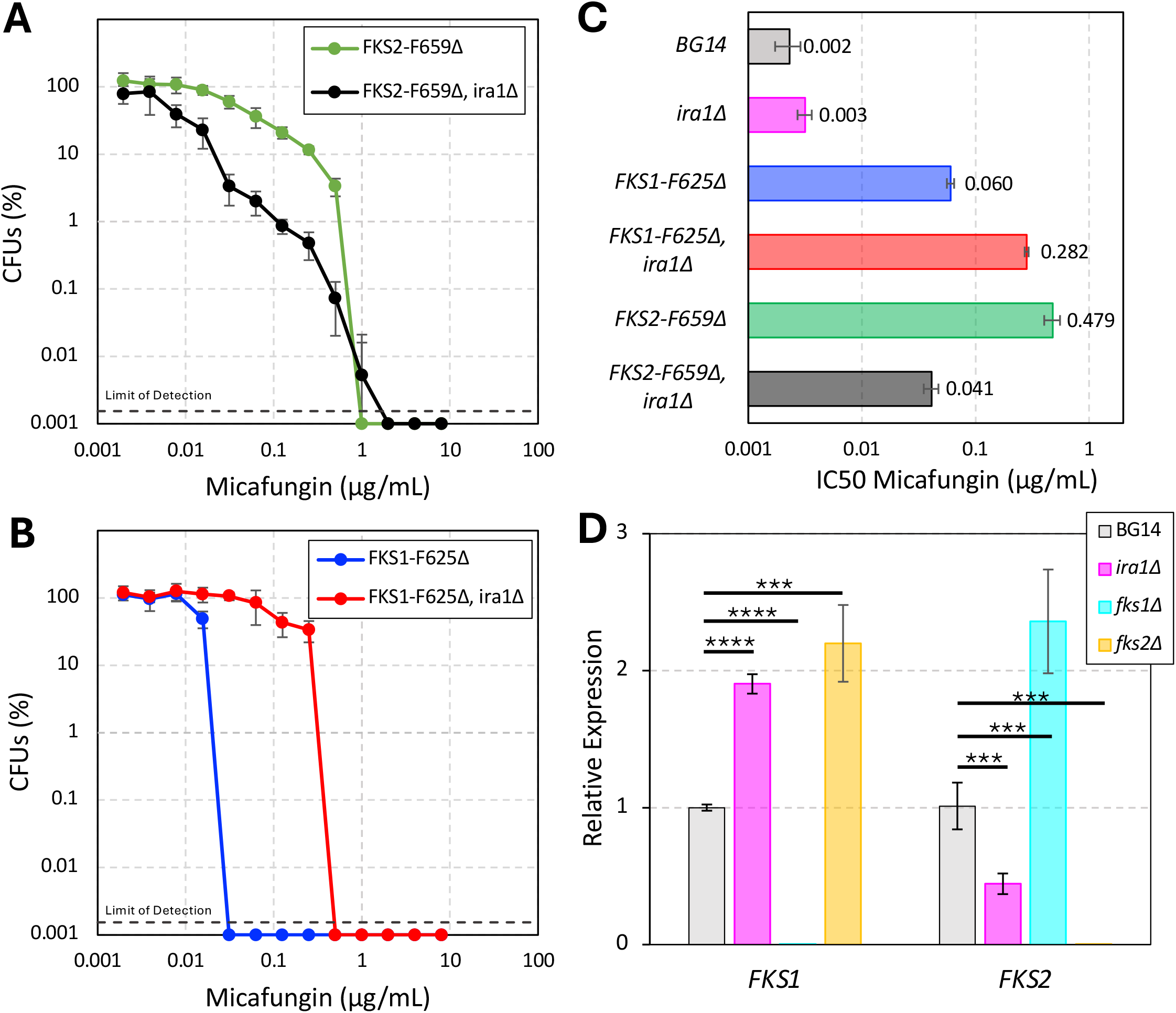
Ira1 regulates heteroresistance and resistance. The *IRA1* gene was knocked out in the *FKS1-F625Δ, FKS2-F659Δ*, and wild-type BG14 parent strain backgrounds, and the resulting strains were subjected to PAP assays using micafungin as described in Fig. 1 (A, B). The strains were also subjected to broth microdilution assays to quantify micafungin resistance (C). Bars indicated average IC50 of four replicates (±SD). (D) qRT-PCR analysis of *FKS1* and *FKS2* gene expression was performed on wild-type, *ira1*Δ, *fks1Δ*, and *fks2Δ* mutant derivatives of strain BG2 (average of 4 replicates ±SD). Statistical significance was assessed using a Welch’s t-test (***P < 0.005; ****P < 0.001).

We then tested whether Ira1 exhibited similar effects on Fks1-F625Δ. Surprisingly, the *FKS1-F625Δ ira1Δ* double mutant exhibited strongly increased resistance to micafungin relative to the parent strain, as indicated in the PAP assay (Fig. 5B) as well as broth microdilution assay (Fig. 5C). The same patterns were observed in broth microdilution assays using caspofungin (Fig. S6). *IRA1* therefore served as a positive regulator of *FKS2-F659Δ* and a negative regulator of *FKS1-F625Δ*, potentially impacting the mutual antagonism as well. To investigate these disparate effects of Ira1, expression of *FKS2* or *FKS1* genes was quantified in unstressed cells by quantitative real-time PCR. Consistent with the PAP assays, *FKS1* expression was nearly doubled and *FKS2* expression was halved in the *ira1Δ* strain, relative to the parent strain (Fig. 5D). *FKS1* and *FKS2* expression was also increased in an *fks2Δ* mutant and an *fks1Δ* mutant, respectively. This indicates that Ira1 plays a role in the differential expression of the Fks enzymes.

Altogether, the findings suggest that broad echinocandin heteroresistance in *C. glabrata* can arise through cell-to-cell variation in the expression of *FKS2-GOR* gene through a combination of stress and glucose signaling pathways (see Fig. 6). Antagonism by drug-bound Fks1 and a lipid code [26] may also impact Fks2-GOR function and expression and regulate echinocandin heteroresistance.

**Figure 6.**
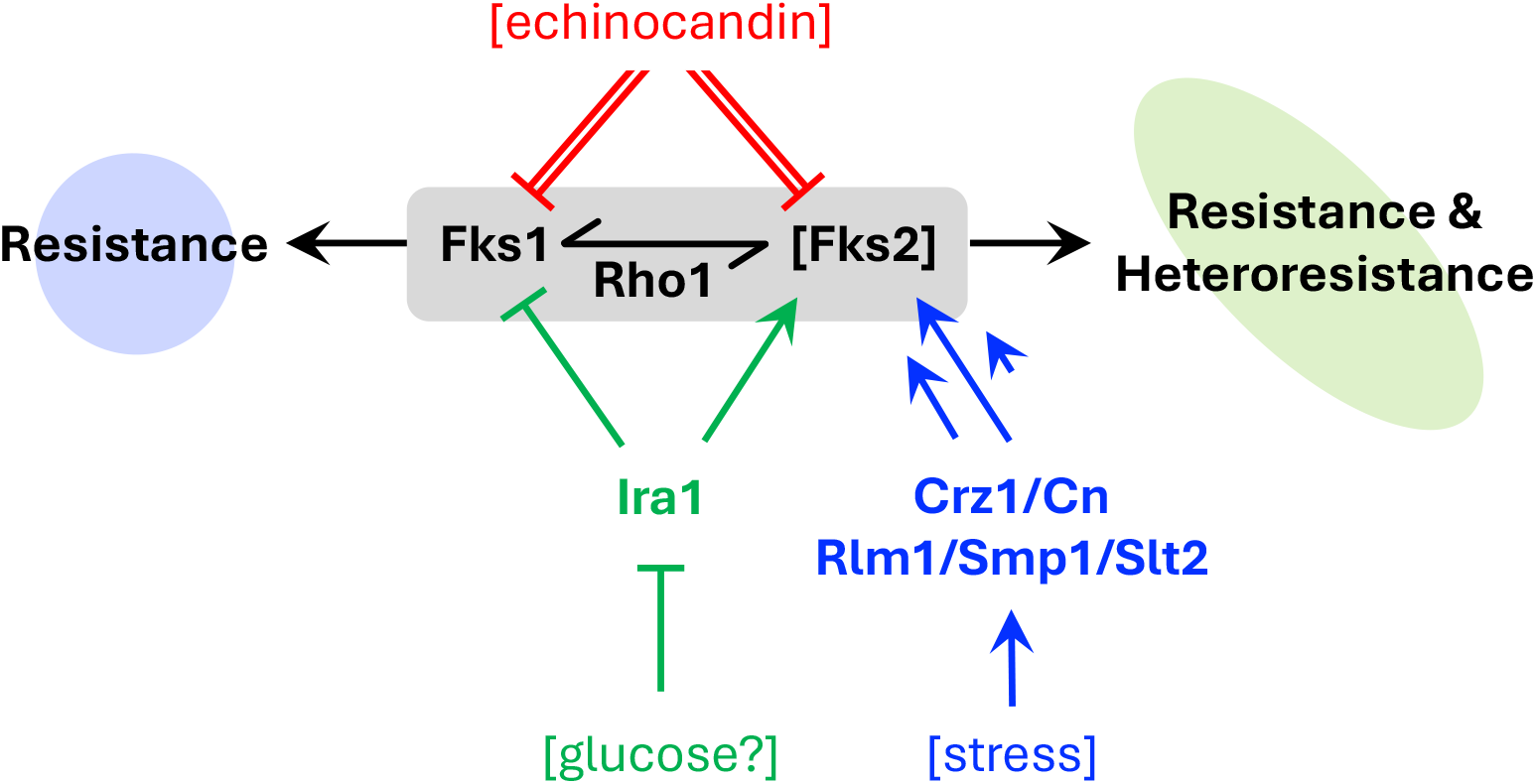
Proposed mechanism of Fks1-Fks2 antagonism and the regulation of heteroresistance and resistance in *C. glabrata*. Echinocandins inhibit wild-type Fks1 and Fks2 enzymes. When one enzyme contains a GOR mutation in hotspot-1 (thus removing a red double arrow), the other remains inhibited at normal doses and antagonizes the function of resistant enzyme by sequestering Rho1 or by another mechanism (see Discussion). This mutual antagonism can be abolished by a secondary loss-of-function or gain-of-resistance mutation in the antagonizing gene (removing the remaining red double arrow and causing super-resistance). Heterogeneous expression of *FKS2-GOR* in clonal populations (indicated by thee arrows of different lengths) is achieved through variation in stress and glucose signaling pathways, resulting in strong heteroresistance resistance. Brackets denote potentially variable components of the system. Ira1 decreases resistance of *FKS1-GOR*, potentially through downstream effectors such as Ras1-cAMP-PKA signaling.

## DISCUSSION

This study reports the first instances of heteroresistance to echinocandins in *C. glabrata*. The underlying regulatory mechanism involved the variable expression of a strong resistance gene, *FKS2-F659Δ*, in clonal populations leading to the appearance of progressive heteroresistance. Multiple lines of evidence support this hypothesis. First, two stress-sensitive transcription factors, Crz1 and Rlm1, were found to be important for maximum CFUs in the transitional range of micafungin doses and their upstream regulators, calcineurin and Slt2, were found to be even more important. Second, activation or inhibition of calcineurin for just 2 hours prior to PAP assays correspondingly increased or decreased CFUs in the transitional range of micafungin, respectively, in *FKS2-F659Δ* mutant populations. Finally, a recent study showed broad cell-to-cell variation in the expression of functional halo-tagged Fks2 protein in wild-type strain backgrounds [10]. Progressive heteroresistance was observed with multiple echinocandins and in two highly divergent strains of *C. glabrata*. Additionally, other gain-of-resistance (GOR) mutations in *FKS2* also exhibited elevated levels of heteroresistance. As GOR mutations in *FKS2* have been found in many patients undergoing treatment with echinocandins, the ensuing heteroresistance may contribute to therapeutic failures.

In contrast to the strong heteroresistance observed for *FKS2-GOR* mutants, the corresponding mutations in *FKS1* produced little or no heteroresistance despite the strong resistance to echinocandins. Expression of *FKS1* appears unresponsive to the stress signaling pathways and more constitutive, possibly producing less variation of Fks1 protein levels from cell-to-cell. Flow-cytometry of cells expressing halo-tagged Fks1 could be used to confirm that prediction [10]. Resistance of *FKS1-F625Δ* mutants to echinocandins was not strongly affected by calcineurin inhibitors either in broth microdilution assays [8] or PAP assays, suggesting that calcineurin signaling did not strongly impact its expression or function. However, the *FKS1-F625Δ* mutants of *C. glabrata* analyzed here did indeed exhibit significant CFUs over a narrow range of transitional micafungin doses, suggesting the acquisition of mild heteroresistance. This mild heteroresistance conferred by constitutively expressed Fks1-F625Δ protein may be due, at least partly, to the antagonistic effects of drug-bound Fks2 that itself is expressed heterogeneously in the population. Resistance of *FKS1-GOR* mutants and *FKS2-GOR* mutants has long been known to be antagonized by the presence of the echinocandin-sensitive paralog [8]. Consequently, “super-resistance” can arise through secondary mutations where the paralog was either destroyed [8, 47] or acquired a secondary GOR mutation [17, 48]. Such super-resistance has been observed also in clinical isolates from patients receiving echinocandin therapies [8]. Based on previous studies in *S. cerevisiae*, at least three different, but non-exclusive, hypotheses could explain the observed antagonism between drug-bound Fks proteins and drug-free Fks-GOR proteins. First, drug-bound Fks proteins could sequester Rho1 and thereby lower activity of the Fks-GOR proteins particularly if Rho1-GTP levels are limiting, which seems to be the case in *S. cerevisiae* [49]. Second, drug-bound Fks proteins could directly sequester the drug-free Fks-GOR proteins into heterodimers with reduced activity. Consistent with this idea, inactive homodimers of Fks1 that are bridged by a tRNA molecule and occluded from Rho1 binding have been isolated recently from *S. cerevisiae* and visualized by cryo-EM [50]. Third, drug-bound Fks proteins may occupy important sites in the cell membrane and prevent access of those sites to the active drug-free Fks-GOR enzymes. The halo-tagged Fks1 and Fks2 proteins appeared to occupy partially overlapping but still distinct domains in the growing buds of otherwise wild-type *C. glabrata* [10]. In any of these scenarios, variable expression of *FKS2* in the population could produce heterogenous levels of antagonism in *FKS1-GOR* mutants and thereby lead to increased heteroresistance. These hypotheses surrounding antagonism are not mutually exclusive of one another and each may contribute significantly to the observed patterns of resistance and heteroresistance in *C. glabrata*.

The mechanisms proposed here may help explain echinocandin heteroresistance and resistance in other *Candida* species. In *C. parapsilosis*, where echinocandin heteroresistance has been observed in many isolates and linked to therapeutic failure [25], a natural gain-of-resistance mutation exists in hotspot-1 of *FKS1* [31]. The progressive heteroresistance observed in some of those isolates could arise from heterogeneous expression of *FKS1-GOR* in clonal populations or from variable levels of antagonism of the drug-free Fks1-GOR by the drug-bound Fks2 or Fks3 proteins, which lack the GOR mutation. The *FKS2* and *FKS3* genes are highly expressed in *C. parapsilosis* [31] and FK506 blocked *C. parapsilosis* growth in laboratory conditions [51], suggesting the species may experience stress in that environment. The non-progressive heteroresistance observed in some isolates of *C. parapsilosis* could arise from unstable aneuploidy or unstable copy number variation or other mechanisms that remain to be discovered. In *C. albicans*, the non-essential *FKS2* and *FKS3* genes have been shown to function as negative regulators of echinocandin resistance, which primarily depends on the essential *FKS1* gene encoding a susceptible enzyme [52]. When gain-of-resistance mutations in *FKS1* arise heterozygously in diploid *C. albicans*, the resistance is initially antagonized by the drug-sensitive allele and super-resistance occurs upon secondary loss-of-function mutations or gain-of-resistance mutations in that allele [53]. Echinocandin heteroresistance in *C. albicans*, which has not yet been reported [24], may only arise when gain-of-resistance mutations in *FKS1* become homozygous (diminishing antagonism by the wild-type *FKS1* allele) and when antagonism by *FKS2* and *FKS3* becomes elevated heterozygously in the population. An instance of echinocandin heteroresistance reported in *C. auris* [25] may be explained by any of these mechanisms.

Our Tn-seq profiling in *C. glabrata* suggests that the gene networks controlling heteroresistance to caspofungin (in *FKS2-F659Δ* mutants) are largely distinct from those controlling resistance to caspofungin (in wild-type cells). Yet, at the highest doses of caspofungin treatment, drug susceptibility was still mediated by sphingolipid genes *FKS2-F659Δ* mutants, as in wild-type cells. One of the strongest hits produced complex effects: *IRA1*, a component of glucose-sensing pathways that regulate cAMP levels and protein kinase A signaling [29], was required to increase heteroresistance and resistance of *FKS2-F659Δ* mutants and limited resistance of *FKS1-F625Δ* mutants via differential transcriptional regulation of the two enzymes. Other hits may act primarily through one *FKS* gene and secondarily on the other by altering the mutual antagonism. Crz1 was shown to be important for heteroresistance; two genes that exhibited high positive Z-scores in the Tn-seq screen were *DFG5* and *FLC2* which function together as proteins in attachment of GPI-anchored proteins to the cell wall and were previously shown to be targets of Crz1 [40]. These and other proteins involved in cell wall and glucan biogenesis, particularly those that are expressed heterogeneously by calcineurin-Crz1, could post-transcriptionally regulate and antagonize Fks enzymes, thus contributing to heteroresistance. *SSK1* and *SSK2*, MAPKKKs in the HOG signaling pathway [45], were elucidated by the Tn-seq screen as potential negative regulators of heteroresistance in the BG14 strain. However, SSK2 is polymorphic and nonfunctional in the CBS138 strain [45] which may indicate that there are natural variations in heteroresostance between strains of *C. glabrata*. Additional Tn-seq screens using *FKS1-F625Δ* mutants and from derived strains that lack antagonism could help distinguish their mechanisms of action.

The mechanisms of echinocandin heteroresistance may be distinct from mechanisms of echinocandin tolerance and persistence, which are starting to be unraveled in *C. glabrata* [23]. Tolerance and persistence mechanisms increase the lifespan of major and minor subpopulations, respectively, of cells exposed to extremely high doses of echinocandins that are far above the MIC [54]. Though brief stress (e.g. manogepix pretreatment) and calcineurin signaling strongly increased tolerance of wild-type *C. glabrata* to echinocandins, the effects were largely independent of *CRZ1* and completely independent of *FKS2* [23]. Calcineurin therefore sits at a critical control point in *C. glabrata* that confers tolerance and persistence to echinocandins in wild-type strains in addition to resistance and heteroresistance in strains bearing *FKS2-GOR* mutations. The findings presented here suggest that short-term administration of tacrolimus (FK506) or other inhibitors of fungal calcineurin could improve clinical outcomes by blocking all these undesirable phenomena associated with antifungal therapies, while potentially avoiding side-effects (immunosuppression, nephrotoxicity, etc.) that arise from long-term use.

Heteroresistance to azole-class antifungals has also been detected in a broad survey *C. glabrata* clinical isolates [22] and in other fungal species [18, 19, 55, 56]. In *Cryptococcus neoformans*, heteroresistance to azoles may be conferred by heterogeneous dosages of an entire chromosome resulting in overexpression of the target (*ERG11*) and a drug efflux pump [18]. In *C. glabrata*, the degree of fluconazole heteroresistance was strongly correlated with the rate of fluconazole efflux and expression of several efflux pumps [22]. The target of fluconazole, Erg11, was also induced by calcineurin signaling in *C. glabrata* [57], potentially indicating another component to the regulation of azole heteroresistance. Azole tolerance, a phenomenon distinct from heteroresistance, and resistance that may contribute to therapeutic failures, has long been known to depend on calcineurin signaling in *C. glabrata, C. albicans*, and several other fungal species [43, 58, 59]. Deeper exploration of how calcineurin signaling achieves all these anti-antifungal effects could spur the development of non-immunosuppressive compounds that augment the efficacy of multiple classes of antifungals.

## METHODS

### *C. glabrata* Strains and Plasmids

All strains and their sources are listed in Table S3. *C. glabrata* strains were derived from BG14, a *ura3Δ* derivative, of wild-type strain BG2 [60]. Individual gene knockouts were constructed using the PRODIGE method [61] where coding sequences were replaced with the coding sequence of *URA3 or HIS3* from *S. cerevisiae* plasmids pRS406 or pRS403, respectively [62]. Colony PCR was used for screening validation. For this study, mutant FKS1 and FKS2 PCR amplified from FKS1-F625Δ and FKS2-F659Δ in 2001-HTU [8] (received from David Perlin) and transformed into strain BG14. All oligonucleotides used tor generating and screening mutants are available in Table S4.

### Antifungals and key chemicals

Micafungin (Cat. #18009) was obtained from Cayman Chemicals, and caspofungin (Cat. #S3073), tactrolimus (FK506) (Cat. #S5003), and manogepix (E1210) (Cat. #S0491) from SelleckChem. 5-Fluoroorotic Acid (5-FoA) was obtained from Zymo Research (Cat. # F9001-5) and ClonNAT (Nourseothricin Sulfate) was obtained from GoldBio (Cat. #N-500-100).

### Tn-seq screen for genes that regulate caspofungin heteroresistance

A new pool of Hermes-NAT1 insertion mutants in *FKS2-F659Δ ura3Δ* strain was generated by transforming the strain with pCU-MET3-Hermes, then inducing transposition in 40 replicate cultures, as described previously [63]. Replicate cultures were pooled and enriched in 5-FOA and NAT, then cryo-preserved for prior to Tn-seq as previously described [63]. The complex pool of insertion mutants was thawed from storage at −80°C, grown to saturation, then diluted into fresh SCD-0 medium containing a range of caspofungin doses, and shaken at 30°C for 48 h [30]. The cells were pelleted, washed, and then genomic DNA was extracted using the Quick-DNA Fungal/Bacterial Miniprep Kit from Zymo Research (Cat. #D6005). gDNA was fragmented by sonication, repaired, ligated to adapters, size-selected, and then insertion sites were PCR amplified as described previously [64]. Single-end reads were sequenced using AVITI 2×75 Sequencing Kit Cloudbreak FS High Output (Cat. # 860-00015) with custom primers specific to the HERMES transposon and the splinkerette to ensure that all reads were from genomic DNA at the transposon insertion site [64].

Samples were basecalled and demultiplexed using Bases2Fastq software (v2.1.0) by element biosciences and CutAdapt was used to identify and trim away adapter sequences in the case of adapter readthrough [65] and aligned to the BG2v1 reference genome [66] using Bowtie2 [67]. Mapped reads with low quality (Q<20) or mismatches at position 1 were discarded and the remainder were tabulated for each site in the genome. The total number of sequence reads in each annotated gene were tabulated and the top site was removed to limit passenger effects [68]. Z-scores were calculated for each gene at every caspofungin does as before [30] (Table S1). Z-scores for each gene were averaged across the four doses. Likewise, Z-scores from a caspofungin resistance screen in Pool-3 of Hermes-NAT1 insertion mutants in strain BG14 were averaged across the two lowest doses for comparison **(Fig 4, Table S1)**.

### Population Analysis Profile (PAP) assays and analyses

Single colonies were grown to saturation overnight in SCD-0 media, shaking at 30°C. Solid YPD media containing varying doses of drug were prepared by adding drug to warmed, liquid media, and allowing plates to solidify at room temperature for at least 24 hours. The saturated cultures were diluted 1:50 into fresh SCD-0 media and then serially diluted 1:5. 2 µL of each dilution was spotted onto agar YPD media containing varying doses of micafungin or caspofungin. Plates were incubated for 24 hours at 30°C and single colonies were counted manually using a dissecting microscope. All strains were tested in biological quadruplicate. Colony forming units per mL and the percentage of CFUs for each replicate were calculated with respect to the no drug control at every drug concentration and the arithmetic mean and standard deviation was calculated at each drug concentration.

### Microbroth Dilution Assay and analyses

Single colonies were grown to saturation overnight in SCD medium, shaking at 30°C. Cells were back diluted 1:2000 in fresh SCD medium with 1:2 serial dilutions of the antifungal in flat-bottom 96-well plates. Treated cells were grown 20-24 hours standing at 30°C, shaken to resuspend, and OD600 was measured using an Accuris Instruments SmartReader 96T. All strains were tested in biological quadruplicate. The raw data were fitted to a sigmoid equation y = m1+(m2-m1)/(1+(x/m3)^m4 in Kaleidagraph (v5.04, Synergy) and the dose at which half-maximal growth was observed (IC50) was calculated independently for each replicate. The arithmetic mean and standard deviation was calculated for each strain.

### RT-qPCR Experiments

Single colonies were picked and grown overnight, shaking at 30°C to saturation For all samples, cells were diluted 1:50 in fresh SCD medium, and harvested by centrifugation (14 k, 60 s). The supernatant was aspirated, and cell pellets were flash frozen in liquid nitrogen and stored at ™80°C until RNA extraction. Total RNA was extracted from cells using a hot acid phenol-chloroform extraction protocol [69]. Cells were lysed in RNA lysis buffer (6 mM NaOAc, 8.4 mM EDTA, 1% SDS) and RNA was purified through two phenol and one chloroform extraction. RNA was precipitated with isopropanol, resuspended in TE buffer, and treated with DNAse (New England Biolabs). 1µg RNA was reverse transcribed using the High-Capacity cDNA Reverse Transcription Kit (Thermo). Real-time PCR was performed using the CFX96 Touch Real-Time PCR Detection System (BioRad) using the ABsolute Blue QPCR Mix SYBR Green Kit (ThermoFisher) with the following parameters: 15 min at 95°C, 40× (15 min at 95°C, 30 min at 58°C, 30 min at 72°C). Target gene transcript levels were normalized to averaged TEF1 transcript levels in each sample, and this ratio from each sample was normalized to that of BG14 cells. Target primers are identified in Table S4.

### Data availability

The authors affirm that all data necessary for confirming the conclusions of the article are present within the article, figures, tables, and repository. Raw sequencing reads used in this study were deposited at the NCBI Sequence Read Archive (SRA) with the BioProject IDxxxx. Caspofungin resistance data [30] were obtained from SRA BioProject ID PRJNA1247003. A tabulation of the chromosomal coordinates and frequency of each mapped transposon insertion site (site count files) and a tabulation of the number of mapped transposon sites that fall within annotated gene boundaries (gene count files) are available upon request.

## ACKNOWLEDGEMENTS

The authors thank Dr. Winston Timp and Jess Hosea for providing access to DNA sequencing instruments and Dr. John Kim for the use of other critical instruments. We are grateful to Drs. David Perlin, Erika Shor, Alejandro de las Peñas, Anne G. Rosenwald, and Ronda Rolfes for providing *C. glabrata* strains. The authors also thank Timothy Nickels, Anna Zaeske, Caroline Moore, and Fijare Plous for contributing materials and support. This research was supported by grants from the National Institutes of Health (T32-GM007231 to the Cell, Molecular, Developmental Biology, and Biophysics doctoral, training program; R01-AI153414 to KWC).

## REFERENCES

1. Gold, J.A.W., et al., Candida glabrata emerges as the most common cause of candidemia: analysis of a large hospital-based database, United States, 2016-2024. Clin Infect Dis, 2026.

2. Guinea, J., Global trends in the distribution of Candida species causing candidemia. Clin Microbiol Infect, 2014. 20 Suppl 6: p. 5–10.

3. Pfaller, M.A., et al., Twenty Years of the SENTRY Antifungal Surveillance Program: Results for Candida Species From 1997-2016. Open Forum Infect Dis, 2019. 6(Suppl 1): p. S79–S94.

4. Beardsley, J., et al., Candida glabrata (Nakaseomyces glabrata): A systematic review of clinical and microbiological data from 2011 to 2021 to inform the World Health Organization Fungal Priority Pathogens List. Med Mycol, 2024. 62(6).

5. Shields, R.K., et al., Five-minute exposure to caspofungin results in prolonged postantifungal effects and eliminates the paradoxical growth of Candida albicans. Antimicrob Agents Chemother, 2011. 55(7): p. 3598–602.

6. Perlin, D.S., Mechanisms of echinocandin antifungal drug resistance. Ann N Y Acad Sci, 2015. 1354(1): p. 1–11.

7. Marcet-Houben, M. and T. Gabaldón, Beyond the Whole-Genome Duplication: Phylogenetic Evidence for an Ancient Interspecies Hybridization in the Baker’s Yeast Lineage. PLoS Biol, 2015. 13(8): p. e1002220.

8. Katiyar, S.K., et al., Fks1 and Fks2 are functionally redundant but differentially regulated in Candida glabrata: implications for echinocandin resistance. Antimicrob Agents Chemother, 2012. 56(12): p. 6304–9.

9. Healey, K.R., et al. Differential Regulation of Echinocandin Targets Fks1 and Fks2 in Candida glabrata by the Post-Transcriptional Regulator Ssd1. Journal of Fungi, 2020. 6, 143 DOI: 10.3390/jof6030143.

10. Gonzalez-Jimenez, I., et al., Expression of 1,3-β-glucan synthase subunits in Candida glabrata is regulated by the cell cycle and growth conditions and at both transcriptional and post-transcriptional levels. Antimicrobial Agents and Chemotherapy, 2025. 69(8): p. e00500–25.

11. Garcia-Effron, G., S. Park, and D.S. Perlin, Correlating echinocandin MIC and kinetic inhibition of fks1 mutant glucan synthases for Candida albicans: implications for interpretive breakpoints. Antimicrob Agents Chemother, 2009. 53(1): p. 112–22.

12. Pham, C.D., et al., Role of FKS Mutations in Candida glabrata: MIC Values, Echinocandin Resistance, and Multidrug Resistance. Antimicrobial Agents and Chemotherapy, 2014. 58(8): p. 4690–4696.

13. Aldejohann Alexander, M., et al., In vitro activity of ibrexafungerp against clinically relevant echinocandin-resistant Candida strains. Antimicrobial Agents and Chemotherapy, 2024. 68(2): p. e01324–23.

14. You, Z.L., et al., Inhibition mechanism of the fungal β-1,3-glucan synthases by triterpenoid antifungal drugs. Nat Commun, 2026. 17(1).

15. Castanheira, M., et al., Frequency of fks mutations among Candida glabrata isolates from a 10-year global collection of bloodstream infection isolates. Antimicrob Agents Chemother, 2014. 58(1): p. 577–80.

16. Alexander, B.D., et al., Increasing echinocandin resistance in Candida glabrata: clinical failure correlates with presence of FKS mutations and elevated minimum inhibitory concentrations. Clin Infect Dis, 2013. 56(12): p. 1724–32.

17. Zajac, C., et al., Hotspot gene conversion between FKS1 and FKS2 in echinocandin resistant Candida glabrata serial isolates. npj Antimicrobials and Resistance, 2025. 3(1): p. 31.

18. Sionov, E., et al., Cryptococcus neoformans Overcomes Stress of Azole Drugs by Formation of Disomy in Specific Multiple Chromosomes. PLOS Pathogens, 2010. 6(4): p. e1000848.

19. Sionov, E., Y.C. Chang, and K.J. Kwon-Chung, Azole Heteroresistance in Cryptococcus neoformans: Emergence of Resistant Clones with Chromosomal Disomy in the Mouse Brain during Fluconazole Treatment. Antimicrobial Agents and Chemotherapy, 2013. 57(10): p. 5127–5130.

20. Huang, Y., et al., Epidemiology of cryptococcal meningitis and fluconazole heteroresistance in Cryptococcus neoformans isolates from a teaching hospital in southwestern China. Microbiol Spectr, 2024. 12(8): p. e0072524.

21. Moreira, I.M.B., et al., Fluconazole Resistance and Heteroresistance in Cryptococcus spp.: Mechanisms and Implications. Rev Soc Bras Med Trop, 2025. 58: p. e002002025.

22. Ben-Ami, R., et al., Heteroresistance to Fluconazole Is a Continuously Distributed Phenotype among Candida glabrata Clinical Strains Associated with In Vivo Persistence. mBio, 2016. 7(4): p. 10.1128/mbio.00655-16.

23. Harrington, A.A., T.J. Nickels, and K.W. Cunningham, Echinocandin tolerance and persistence in vitro are regulated by calcineurin signaling in Candida glabrata. mBio, 2026. 17(1): p. e0254625.

24. Shor, E., S. Perlin David, and P. Kontoyiannis Dimitrios, Tolerance and heteroresistance to echinocandins in Candida auris: conceptual issues, clinical implications, and outstanding questions. mSphere, 2025. 10(5): p. e00161–25.

25. Zhai, B., et al., Antifungal heteroresistance causes prophylaxis failure and facilitates breakthrough Candida parapsilosis infections. Nat Med, 2024. 30(11): p. 3163–3172.

26. El-Halfawy, O.M. and M.A. Valvano, Antimicrobial Heteroresistance: an Emerging Field in Need of Clarity. Clinical Microbiology Reviews, 2015. 28(1): p. 191–207.

27. Gautier, C., I. Maciel Eli, and V. Ene Iuliana, Approaches for identifying and measuring heteroresistance in azole-susceptible Candida isolates. Microbiology Spectrum, 2024. 12(4): p. e04041–23.

28. Lyons, N. and J. Berman, Protocols for Measuring Tolerant and Heteroresistant Drug Responses of Pathogenic Yeasts, in Antifungal Drug Resistance: Methods and Protocols, D.J. Krysan and W.S. Moye-Rowley, Editors. 2023, Springer US: New York, NY. p. 67–79.

29. Tanaka, K., K. Matsumoto, and E.A. Toh, IRA1, an inhibitory regulator of the RAS-cyclic AMP pathway in Saccharomyces cerevisiae. Mol Cell Biol, 1989. 9(2): p. 757–68.

30. Nickels Timothy, J. and W. Cunningham Kyle, Tn-seq screens in Candida glabrata treated with echinocandins and ibrexafungerp reveal pathways of antifungal resistance and cross-resistance. mSphere, 2025. 10(7): p. e00270–25.

31. Garcia-Effron, G., et al., A naturally occurring proline-to-alanine amino acid change in Fks1p in Candida parapsilosis, Candida orthopsilosis, and Candida metapsilosis accounts for reduced echinocandin susceptibility. Antimicrob Agents Chemother, 2008. 52(7): p. 2305–12.

32. Sherman, E.X., J.E. Wozniak, and D.S. Weiss, Methods to Evaluate Colistin Heteroresistance in Acinetobacter baumannii. Methods Mol Biol, 2019. 1946: p. 39–50.

33. Kitada, K., E. Yamaguchi, and M. Arisawa, Cloning of the Candida glabrata TRP1 and HIS3 genes, and construction of their disruptant strains by sequential integrative transformation. Gene, 1995. 165(2): p. 203–6.

34. Schwarzmuller, T., et al., Systematic phenotyping of a large-scale Candida glabrata deletion collection reveals novel antifungal tolerance genes. PLoS Pathog, 2014. 10(6): p. e1004211.

35. Nickels, T.J., et al., Transposon-sequencing (Tn-seq) of the Candida glabrata reference strain CBS138 reveals epigenetic plasticity, structural variation, and intrinsic mechanisms of resistance to micafungin. G3 Genes|Genomes|Genetics, 2024. 14(9): p. jkae173.

36. Pfaller, M.A., et al., Frequency of decreased susceptibility and resistance to echinocandins among fluconazole-resistant bloodstream isolates of Candida glabrata. J Clin Microbiol, 2012. 50(4): p. 1199–203.

37. Misas, E., et al., Genomic description of acquired fluconazole- and echinocandin-resistance in patients with serial Candida glabrata isolates. Journal of Clinical Microbiology, 2024. 62(2): p. e01140–23.

38. Zimbeck Alicia, J., et al., FKS Mutations and Elevated Echinocandin MIC Values among Candida glabrata Isolates from U.S. Population-Based Surveillance. Antimicrobial Agents and Chemotherapy, 2010. 54(12): p. 5042–5047.

39. Levin, B.R., et al., Theoretical considerations and empirical predictions of the pharmaco- and population dynamics of heteroresistance. Proc Natl Acad Sci U S A, 2024. 121(16): p. e2318600121.

40. Pavesic Matthew, W., et al., Calcineurin-dependent contributions to fitness in the opportunistic pathogen Candida glabrata. mSphere, 2024. 9(1): p. e00554–23.

41. Singh-Babak, S.D., et al., Global Analysis of the Evolution and Mechanism of Echinocandin Resistance in Candida glabrata. PLOS Pathogens, 2012. 8(5): p. e1002718.

42. Zhao, C., et al., Temperature-induced expression of yeast FKS2 is under the dual control of protein kinase C and calcineurin. Mol Cell Biol, 1998. 18(2): p. 1013–22.

43. Bonilla, M., K.K. Nastase, and K.W. Cunningham, Essential role of calcineurin in response to endoplasmic reticulum stress. Embo j, 2002. 21(10): p. 2343–53.

44. Dudgeon, D.D., et al., Nonapoptotic death of Saccharomyces cerevisiae cells that is stimulated by Hsp90 and inhibited by calcineurin and Cmk2 in response to endoplasmic reticulum stresses. Eukaryot Cell, 2008. 7(12): p. 2037–51.

45. Gregori, C., et al., The High-Osmolarity Glycerol Response Pathway in the Human Fungal Pathogen Candida glabrata Strain ATCC 2001 Lacks a Signaling Branch That Operates in Baker’s Yeast. Eukaryotic Cell, 2007. 6(9): p. 1635–1645.

46. Eden, E., et al., GOrilla: a tool for discovery and visualization of enriched GO terms in ranked gene lists. BMC Bioinformatics, 2009. 10(1): p. 48.

47. Ksiezopolska, E., et al., Narrow mutational signatures drive acquisition of multidrug resistance in the fungal pathogen Candida glabrata. Curr Biol, 2021. 31(23): p. 5314–5326.e10.

48. Saraya, T., et al., Breakthrough invasive Candida glabrata in patients on micafungin: a novel FKS gene conversion correlated with sequential elevation of MIC. J Clin Microbiol, 2014. 52(7): p. 2709–12.

49. Xu, S., et al., Effects of Rho1, a small GTPase on the production of recombinant glycoproteins in Saccharomyces cerevisiae. Microb Cell Fact, 2016. 15(1): p. 179.

50. Li, J., et al., Structural-guided identification of two modulators of β™1,3-glucan synthase FKS1. Nature Communications, 2025. 17(1): p. 591.

51. Cho, O., et al., Tacrolimus (FK506) Exhibits Fungicidal Effects against Candida parapsilosis Sensu Stricto via Inducing Apoptosis. J Fungi (Basel), 2023. 9(7).

52. Suwunnakorn, S., et al., FKS2 and FKS3 Genes of Opportunistic Human Pathogen Candida albicans Influence Echinocandin Susceptibility. Antimicrob Agents Chemother, 2018. 62(4).

53. Douglas, C.M., et al., Identification of the FKS1 gene of Candida albicans as the essential target of 1,3-beta-D-glucan synthase inhibitors. Antimicrob Agents Chemother, 1997. 41(11): p. 2471–9.

54. Brauner, A., et al., Distinguishing between resistance, tolerance and persistence to antibiotic treatment. Nature Reviews Microbiology, 2016. 14(5): p. 320–330.

55. Escribano, P., et al., In Vitro Acquisition of Secondary Azole Resistance in Aspergillus fumigatus Isolates after Prolonged Exposure to Itraconazole: Presence of Heteroresistant Populations. Antimicrobial Agents and Chemotherapy, 2012. 56(1): p. 174–178.

56. Varma, A. and K.J. Kwon-Chung, Heteroresistance of Cryptococcus gattii to Fluconazole. Antimicrobial Agents and Chemotherapy, 2010. 54(6): p. 2303–2311.

57. Wang, Q., et al., Two Sequential Clinical Isolates of Candida glabrata with Multidrug-Resistance to Posaconazole and Echinocandins. Antibiotics (Basel), 2021. 10(10).

58. Marchetti, O., et al., Fungicidal synergism of fluconazole and cyclosporine in Candida albicans is not dependent on multidrug efflux transporters encoded by the CDR1, CDR2, CaMDR1, and FLU1 genes. Antimicrob Agents Chemother, 2003. 47(5): p. 1565–70.

59. Onyewu, C., et al., Ergosterol biosynthesis inhibitors become fungicidal when combined with calcineurin inhibitors against Candida albicans, Candida glabrata, and Candida krusei. Antimicrob Agents Chemother, 2003. 47(3): p. 956–64.

60. Cormack, B.P. and S. Falkow, Efficient Homologous and Illegitimate Recombination in the Opportunistic Yeast Pathogen Candida glabrata. Genetics, 1999. 151(3): p. 979–987.

61. Edlind, T.D., et al., Promoter-dependent disruption of genes: simple, rapid, and specific PCR-based method with application to three different yeast. Current Genetics, 2005. 48(2): p. 117–125.

62. Sikorski, R.S. and P. Hieter, A system of shuttle vectors and yeast host strains designed for efficient manipulation of DNA in Saccharomyces cerevisiae. Genetics, 1989. 122(1): p. 19–27.

63. Gale, A.N., et al., Identification of Essential Genes and Fluconazole Susceptibility Genes in Candida glabrata by Profiling Hermes Transposon Insertions. G3 (Bethesda), 2020. 10(10): p. 3859–3870.

64. Anna M Zaeske, A.A.H., Timothy J Nickels, Andrew N Gale, Dr. Winston Timp, Nicole Alayo, Yasmine Hassoun, Prof. David S Perlin, Dr. Erika Shor, Dr. Kyle W Cunningham, Facile generation of Hermes insertion mutants in prototrophic Candida glabrata for use in nutrient-limited environments. Microbiology Spectrum, 2026 (Submitted).

65. Martin, M., CUTADAPT removes adapter sequences from high-throughput sequencing reads. EMBnet.journal, 2011. 17.

66. Xu, Z., et al., Cell wall protein variation, break-induced replication, and subtelomere dynamics in Candida glabrata. Molecular Microbiology, 2021. 116(1): p. 260–276.

67. Langmead, B. and S.L. Salzberg, Fast gapped-read alignment with Bowtie 2. Nature Methods, 2012. 9(4): p. 357–359.

68. Michel, A.H., et al., Functional mapping of yeast genomes by saturated transposition. eLife, 2017. 6: p. e23570.

69. Guydosh, Nicholas R. and R. Green, Dom34 Rescues Ribosomes in 3′ Untranslated Regions. Cell, 2014. 156(5): p. 950–962.

